# BATL: Bayesian annotations for targeted lipidomics

**DOI:** 10.1101/2021.03.18.435788

**Authors:** Justin G. Chitpin, Anuradha Surendra, Thao T. Nguyen, Graeme P. Taylor, Hongbin Xu, Irina Alecu, Roberto Ortega, Julianna J. Tomlinson, Angela M. Crawley, Michaeline McGuinty, Michael G. Schlossmacher, Rachel Saunders-Pullman, Miroslava Cuperlovic-Culf, Steffany A.L. Bennett, Theodore J. Perkins

## Abstract

**Motivation:** Bioinformatic tools capable of annotating, rapidly and reproducibly, large, targeted lipidomic datasets are limited. Specifically, few programs enable high-throughput peak assessment of liquid chromatography-electrospray ionization tandem mass spectrometry (LC-ESI-MS/MS) data acquired in either selected or multiple reaction monitoring (SRM and MRM) modes.

**Results:** We present here Bayesian Annotations for Targeted Lipidomics (BATL), a Gaussian naïve Bayes classifier for targeted lipidomics that annotates peak identities according to eight features related to retention time, intensity, and peak shape. Lipid identification is achieved by modelling distributions of these eight input features across biological conditions and maximizing the joint posterior probabilities of all peak identities at a given transition. When applied to sphingolipid and glycerophosphocholine SRM datasets, we demonstrate over 95% of all peaks are rapidly and correctly identified.

**Availability and implementation:** BATL software is freely accessible online at https://complimet.ca/batl/ and is compatible with Safari, Firefox, Chrome and Edge.

**Supplementary information:** Supplementary data are available at *Bioinformatics* online.

## 1. Introduction

Targeted lipidomics employs liquid chromatography coupled to tandem mass spectrometry via electrospray ionization (LC-ESI-MS/MS). Using selected and multiple reaction monitoring (SRM and MRM) modes, pairs of precursor and product ions (transitions), are monitored to quantify lipids of interest. Targeted transition lists are constructed based on prior knowledge of lipid fragmentation pathways as reported in literature, (e.g., (Murphy and Axelsen, 2011)), obtained through exploration of MS/MS spectra for untargeted lipidomic analyses, and/or by performing other semi-targeted, unbiased lipid approaches, such as prior assessment of a given matrix in precursor ion scan mode (Sartain, et al., 2011). Once precursor and product ion pairs are identified, parking on a single product ion effectively reduces interfering signals generated by isobaric lipids from other classes, enabling SRM and MRM modes to excel at high-throughput quantitation of both high and low abundance species (Bowden, et al., 2017). Conversely, high-resolution mass spectrometry-based targeted lipidomics is done by parallel reaction monitoring (PRM). This method exploits instrument setups that combine a quadrupole, a high-energy collisional dissociation (HCD) cell, and a high-resolution mass analyzer such as an Orbitrap or time-of-flight (ToF). The targeted precursor ion is isolated by the quadrupole, fragmented by the HCD, and product ions are simultaneously monitored and quantified by either an Orbitrap or ToF mass analyzer (Gallien, et al., 2014; Peterson, et al., 2012). Parallel monitoring of all product ions eliminates the needs for *a priori* targeted transition lists. Because the precursor ion is selected by the low-resolution quadrupole, this targeted approach remains subject to isobaric contamination and thus requires additional bioinformatic tools to confirm peak identities. Together, these approaches have been used to successfully map fluid and cell-specific lipidomes (Quehenberger, et al., 2010; Sartain, et al., 2011; Slatter, et al., 2016), reveal lipidomic disruptions across biological conditions (Alecu and Bennett, 2019; Wang, et al., 2018), and predict changes in lipid metabolism associated with disease progression (Alshehry, et al., 2016; Blasco, et al., 2017; Granger, et al., 2019).

Despite the power of SRM, MRM, and PRM approaches to quantify lipid analytes, it remains challenging to annotate lipid identities rapidly and reproducibly across large numbers of MS chromatograms, notably when collected from different organisms or matrices using different mass spectrometry methodologies. While the concept of targeting individual lipid species in SRM, MRM, and PRM modes appears straightforward, ensuring peaks are correctly assigned is labour-intensive and not trivial, as exemplified in Figure 1. Multiple isobars, isomers, and isotopologues, sharing the same product ion, can elute in close proximity to the targeted lipid. Moreover, routine variations in chromatography can cause retention time shifts that align isobars or isotopologues with the species of interest in different MS runs (Smith et al., 2015). When multiple peaks are detected at a given transition, careful judgement is required to discriminate between lipid targets. These problems are magnified when researchers seek to match corresponding peaks and identify unique lipid species (a) across lipidomes of different organisms or (b) within different matrices where peak features may change drastically.

**Fig. 1.**
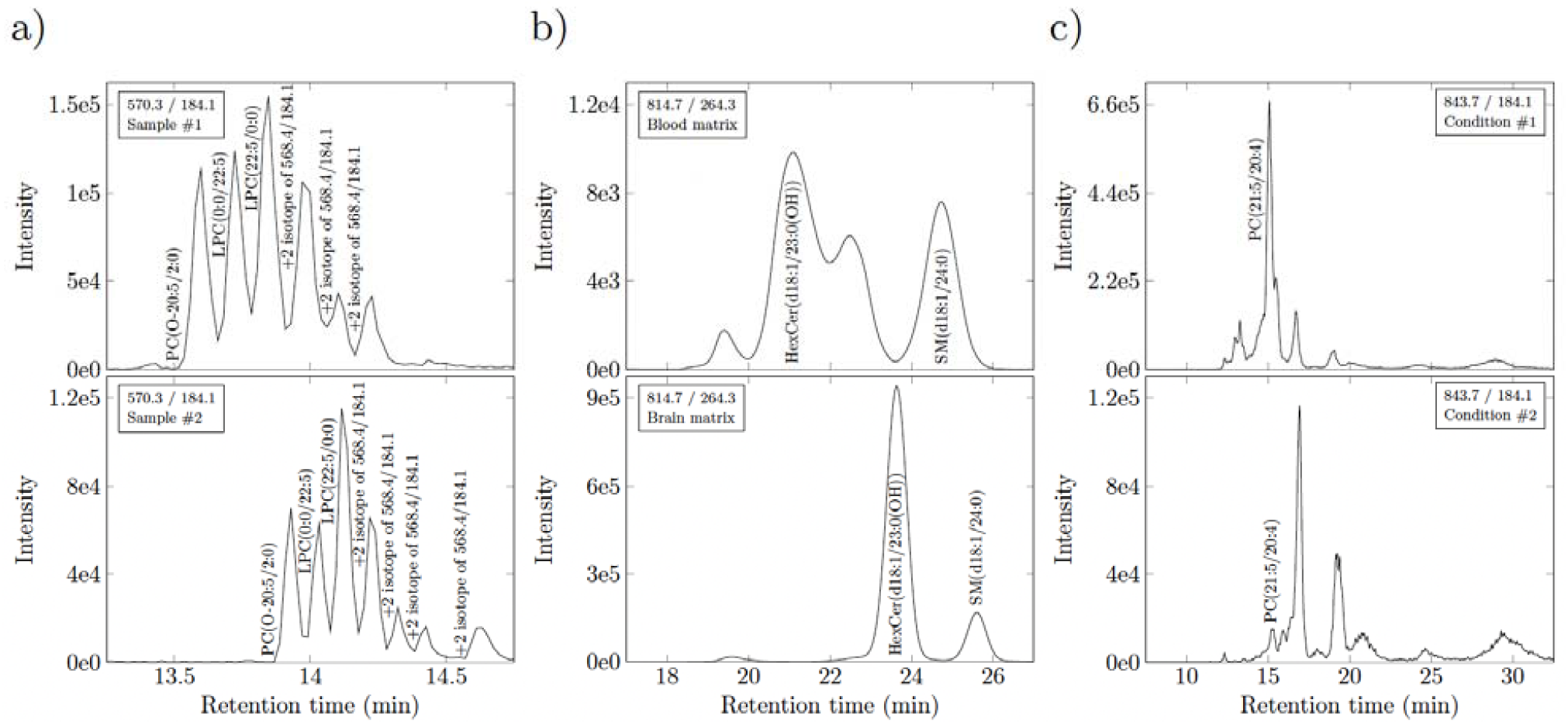
Common challenges associated with SRM, MRM, and PRM peak identification. a) Ambiguity occurs when multiple lipid isomers, isobars, and isotopes are detected within the same matrix at a given transition, yet technical variations in flow rate, composition of the mobile phase, temperature, pH, etc., cause their retention times to vary across samples. Data represent extracted ion chromatograms (XICs) of the same matrix (murine plasma) in animals fed different diets. Note six peaks are observed in one sample at a given transition. Seven peaks are observed in a different sample shifted by one minute. Matching retention time would not align these shifted species. b) Assigning lipid identities based on peak elution order (picking the nth eluting peak) will also lead to misidentifications when comparing lipid species across matrices. Data represent XICs of plasma and brain (temporal cortex) lipidomes from the same animal. Note both the retention time shift and the fundamentally different number of species within each matrix. Matching by either retention time or peak elution order would confound identification. c) Matching lipids based on peak intensity features is complicated by pathological changes detected in lipid metabolism. Data represent XICs of the human plasma lipidome of patients with different neurodegenerative diseases. Note the marked change in abundances between conditions that impacts on lipid identification. While algorithms exist to address each of these challenges, few are applicable to datasets wherein all differences manifest simultaneously. BATL addresses these challenges.

Few programs have been developed to address the difficulties of SRM, MRM, and PRM peak identification. MRMPROBS is the most well-recognized SRM/MRM peak identification program, using a multivariate logistic regression classifier to assign annotations from a library of lipid species (Tsugawa, et al., 2013). The program computes the posterior probability of a peak belonging to a lipid in the training set conditioned on five peak features describing lipid retention time, intensity, and shape. However, these features are reduced to only retention time when classifying SRM peaks. Two further program restrictions of MRMPROS lie in the fact that the number of lipid identities in the training set cannot exceed the number of transitions acquired in the raw MS data and that the compound names in the training set must match the lipid target names in the SRM or MRM method. These restrictions become problematic when new lipid species are discovered in different biological matrices or conditions and users seek to match corresponding lipids across these datasets. mProphet uses a conceptually similar linear discriminant analysis method to identify peptides from SRM and MRM data but further includes addition of decoy transitions that act as negative controls to parameterize the null model and derive false discovery rates (FDRs). These additions improve identification confidence (Reiter, et al., 2011). However, identifying a sufficient number of decoy transitions universally applicable to all lipidomes has proven difficult. Vendor-specific programs such as MultiQuant (SCIEX), MassHunter (Agilent), MassLynx (Waters) and LipidSearch (Thermo Fisher Scientific) are peak-picking algorithms where users can specify retention time windows and compute retention time ratios based on predetermined internal standards to assist in peak identification. MultiQuant, MassHunter, and MassLynx do not, however, assign peak identities. LipidSearch (Thermo Fisher Scientific) assigns peak identity to the closest matching retention time within a user-defined retention time window to a proprietary internal library. Similarly, Lipidyzer, using the Lipidomics Workflow Manager program (SCIEX), assigns lipid identities from differential mobility spectrometry (DMS) data acquired by direct infusion SRM mode (Ubhi, et al., 2016). Additionally, Lipidyzer was designed to analyze data acquired specifically from SCIEX QTRAP 5500/5600 mass spectrometers with a SelexION DMS cell. However, even when using these software packages, manual curation remains the most common peak identification method when extracted ion chromatograms (XICs) do not match exactly to reference samples (Bowden, et al., 2018). Finally, academic programs such as METLIN-MRM (Domingo-Almenara, et al., 2018) use a similar approach to LipidSearch, first aligning XIC peaks by retention time before assigning lipid identities to the closest peak within the retention time window. While these approaches excel in identifying compounds within the same condition in simple matrices, any of the common scenarios described in Figure 1 can lead to peak misidentification.

To address this problem, we applied a Bayesian annotation approach tailored to annotate targeted lipidomic datasets and present the program BATL which overcomes many of the limitations of the manual or template-based curation approaches. BATL is an R package, implemented through an online GUI at CompLiMet: Computational Lipidomics and Metabolomics http://complimet.ca/batl/. The input format is based on results tables generated using MultiQuant (SCIEX) but is applicable to any targeted lipidomics data collection mode from any LC-ESI-MS/MS platform once the user formats their results tables to match the format provided. The program models lipid-specific peak features obtained from a user-curated training set using Gaussian distributions and computes the joint posterior probability of all peak identities in a given sample. BATL was developed using eight specific features describing peak retention, intensity, and shape and the online version allows users to train on any combination of features. We show here that our approach accurately identifies over 95% of all sphingolipid and glycerophosphocholine peaks in SRM datasets analyzed across matrices and disease conditions. Thus, BATL is a useful tool for accurate, targeted lipid identification and, with online access, is easily integrated into any lipidomic pipeline.

## 2. Methods

### 2.1 Overview of program

The BATL workflow is presented in Figure 2. First, a training set is constructed from user-labelled, targeted lipidomic datasets. Second, BATL uses both the training set and the specified input features to construct a naïve Bayes statistical model. Third, the model and associated metadata are exported and used by BATL to annotate peaks in query SRM, MRM, or PRM datasets. If a peak cannot be assigned to an identity present in the training set, an annotation of ‘unassigned’ is returned, enabling the user to assess and validate a potentially novel peak at that transition. An optional BATL function is further included which annotates isotopes in all lipid categories as well as sphingolipid-specific artifacts (e.g. dehydrations, deglycosylations, dimers).

**Fig. 2.**
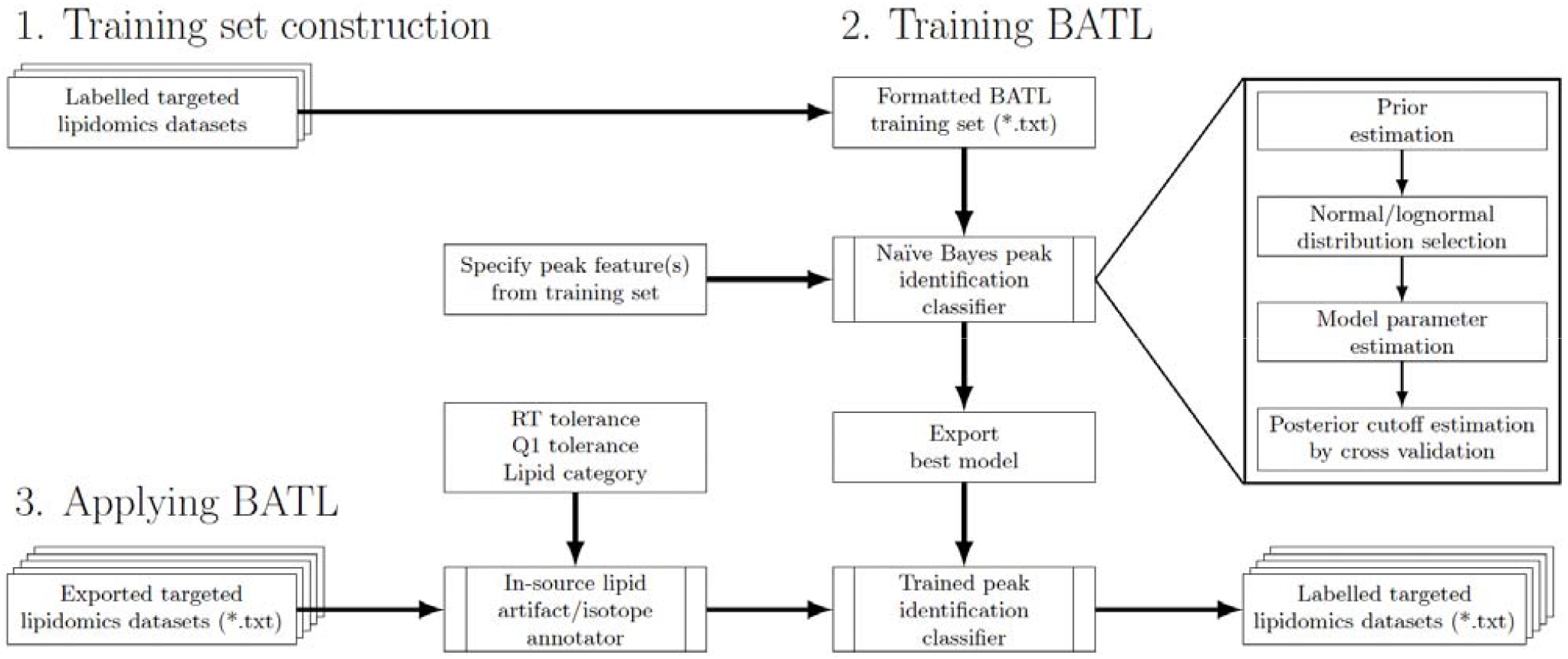
Schematic of the BATL lipid identification workflow. BATL follows three steps: 1) Users are asked to identify training datasets for which they have unambiguous knowledge of peak identities. 2) These datasets are used to train BATL, constructing a naïve Bayes statistical model based on the peak features users select. 3) The model and associated metadata are used by the BATL algorithm to annotate peaks in subsequent query SRM, MRM, or PRM datasets.

### 2.2 Naïve Bayes model

Our approach to peak identification is based on maximizing the joint posterior probability of all peak identities within each sample. Let *P*_1_, *P*_2_,…, *P*_m_ be a list of peaks within a sample described by feature vectors *F*_1_, *F*_2_,…, *F*_*m*_. Each feature vector contains *k* features, where *F*_*i*_ = (*f*_*i*1_, *f*_*i*2_ …, *f*_*ik*_) describes the *i*^th^ peak. Each peak is detected at precursor ion *m*_*i*_ and product ion *p*_*i*_ under the same Q1 and Q3 mass analyzer tolerance δ. Let *I* = {*B*_1_, *B*_2_, …, *B*_*n*_} be the set of all lipid identities, where the *b*^th^ identity is detected at precursor ion *n*_*b*_ and product ion *q*_*b*_ under the Q1 and Q3 mass analyzer tolerance δ. Thus, the possible lipid identities for each peak are those detected within the machine tolerance of the lipid identity and peak transition.

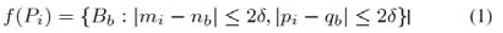

To denote the assigned identity for *P*_*i*_, let *I*_1_, *I*_2_, …, *I*_*m*_ take lipid identities drawn from *f* (*P*_1_), *f* (*P*_2_), …, *f* (*P*_*m*_). The posterior probability of some joint assignment of peak identities is

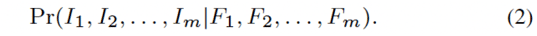

This joint probability is expanded using Bayes’s Theorem as in Equation 3.

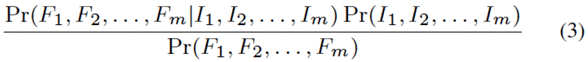

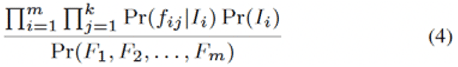

To compute this joint probability, we make three assumptions: 1) the prior probabilities of all lipid identities are independent; 2) the peak feature vectors are statistically independent, conditional on the identities; and 3) the individual features within each vector are statistically independent, conditional on the peak identity. Thus, Equation 3 is simplified to the following probability.

The denominator is a data-dependent constant and can be ignored when comparing the probabilities of different joint assignments. The logposterior probability of a joint assignment is thus proportional to where weight *w*_*ib*_ is the unnormalized, log posterior probability of assigning peak *i* to lipid identity *B*_*b*_.

The joint assignment of lipid identities is determined by the classifier decision rule. To optimize BATL, we tested three classifier rules. First,

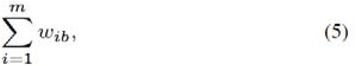

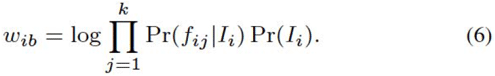

we assessed choosing lipid identities that maximize *w*_*ib*_ following the maximum a posteriori (MAP) decision rule typical of naïve Bayes classifiers. We found that a disadvantage of this decision rule was that lipid identities were assigned independently. Although peaks detected in the same sample clearly corresponded to unique lipid identities, the MAP decision rule could assign an identity more than once per sample (see Results). To address this problem, we evaluated a constrained MAP decision rule wherein lipid identities were assigned by the ranked order of their log posteriors, such that no lipid identity was assigned more than once per sample. We found that this method was not guaranteed to maximize Equation 5 and thus did not yield the optimal assignment of lipid identities (see Results). Third, we resolved the shortcomings of MAP and constrained MAP with the maximum weighted bipartite matching (MWBM) decision rule which considers the simultaneous identification of all peak identities within a sample under the naïve Bayes model.

For every sample transition, a bipartite graph was constructed where the vertices represent peaks *P*_*i*_ and their possible lipid identities *f*(*P*_*i*_) with corresponding edges weighted by *w*_*ib*_. The optimal set of matching peaks and lipid identities was then solved by MWBM, thereby maximizing Equation 5 while ensuring a unique lipid identity was assigned to each peak detected per sample. Finally, under certain conditions, the true identity of a peak would be absent from set *I*, representing a novel lipid species detected in the sample of interest. To account for this possibility, every peak was matched to an ‘unassigned’ identity *U* in addition to *f*(*P*_*i*_). The weights *w*_*iu*_ were found to be specific to each transition and estimated by cross validation (see Results).

### 2.3 Training the model

Let *D* = {*D*_1_, *D*_2_, …, *D*_*p*_} denote the labelled training set containing the instances *D*_*o*_ = (*F*_*o*_, *B*_*o*_) for samples *n* = 1, …, *N*. *F*_*o*_ is the feature vector of length *k*, where *F*_*o*_ = (*f*_*o*1_, *f*_*o*2_, …, *f*_*ok*_), and *B*_*o*_ is the true lipid identity. Each lipid identity in the training set contains a unique sample index because the same lipid can only be detected once per sample. The prior probability of each lipid identity is computed by maximum likelihood estimation

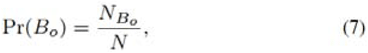

where *N*_*Bo*_ is the number of lipid identities *B*_*o*_ in the training set. The feature likelihoods are computed using either a normal or lognormal distribution with parameters *µ*_*o*_ and *σ*^2^ estimated using the sample mean and variance from the training set. The choice of distribution is assessed using a KS-test for normality and lognormality of feature *j* for lipid identity *B*_o_.

*N*_*j*_ and 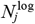 are the number of lipid identities failing the KS-test for normality and lognormality, respectively, for feature *j* at a *P* -value threshold of 0.05.

Lastly, the unassigned identity weights *w*_*iu*_ are estimated per transition by *k*-fold cross validation. Looping over the *k −* 1 folds of the training set, the naïve Bayes model is trained and unnormalized log posteriors are

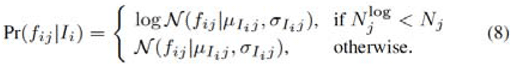

computed from the testing fold. Across all *k* iterations, the weights *w*_*iu*_ for each transition are set to the minimum unnormalized posterior of a correct peak assignment.

### 2.4 Datasets

To train and test BATL, we curated and labelled sphingolipid and glycerophosphocholine datasets composed of 1008 MS spectra generated at the India Taylor Neurolipidomics Research Platform, University of Ottawa. To ensure all of the challenges in MRM, SRM, and PRM identification outlined in Figure 1 were recapitulated in these datasets, we used: (1) a population-based study of circulating lipids in human plasma of cognitively normal controls, and patients suffering from Alzheimer’s Disease, Mild Cognitive Impairment, Dementia with Lewy Bodies, or Parkinson’s Disease (n=319 sphingolipid analyses; n=319 glycerophosphocholine analyses), (2) a genotype and intervention comparison study of lipid metabolism in the temporal cortex, hippocampus, and plasma of wildtype and N5 TgCRND8 mice, a sexually dimorphic mouse model of Alzheimer’s Disease (Granger et al., 2016) (n=121 sphingolipid analyses; n=180 glycerophosphocholine analyses), (3) a technical replicate study of two human plasma samples assessed in 33 sequential runs separated by blanks (n=33 sphingolipid analyses), and as sample data provided online (4) two datasets of human plasma of persons positive or negative for SARS-CoV-2 (n=24 glycerophosphocholine longitudinal analyses provided as two datasets) and (5) a test dataset of human plasma of persons positive or negative for SARS-CoV-2 (n=12 glycerophosphocholine analyses).

To identify all lipids unambiguously in datasets 1-3, all molecular identities were confirmed by LC-SRM-information dependent acquisition (IDA)-enhanced product ion (EPI) experiments of samples pooled across all datasets in which the SRM was used as a survey scan to identify target analytes and an IDA of an EPI spectra was acquired in the linear ion trap and examined to confirm molecular identities. For datasets 4 and 5, each lipid identity was confirmed by LC-IDA-EPI-ESI-MS/MS using SRM as the survey scan. These structural analyses of EPI spectra were further validated by analyzing each lipid (for which commercial standards existed) individually as a standard. All lipids within the sphingolipid dataset were monitored at the same product ion m/z of 264.3 detecting sphingolipids with a d18:1 sphingoid base backbone (sphingosine). All lipids within the glycerophosphocholine dataset were monitored at the same production m/z of 184.1 detecting glycerophospholipids and sphingomyelins with a phosphocholine headgroup. Samples from both the sphingolipid and glycerophosphocholine datasets were equally stratified by acquisition date into training sets for cross validation and holdout sets for model validation. Complete LC-ESI-MS/MS details are provided in Supplementary Data.

### 2.5 Performance metrics

Classifier performance was assessed using metrics of accuracy, identification rate, and unassignment rate. These metrics evaluated how well BATL assigned lipid identities and the calibration of the unassigned identity weights. A correct peak assignment (true positive or TP) was defined as occurring when the classifier assigned the same identity established by IDA-EPI analysis. An incorrect peak assignment (false positive or FP) was defined when the classifier assigned a different identity than the one determined by IDA-EPI structural validation. An unassigned peak (U) refers to when the classifier assigned no identity to the peak (unassigned). Any unassigned peaks were considered incorrectly unassigned when the true identity of all peaks was present in the annotated training set.

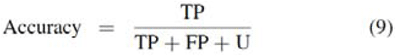

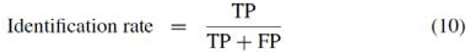

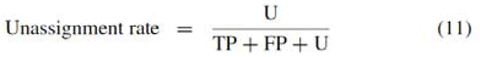

### 2.6 Availability and Implementation

To facilitate use of BATL we have developed a user friendly R/Shiny (Chang, et al., 2021) Web application that enables labelling of MultiQuant, SCIEX data utilizing user and BATL-labelled training datasets. The application with user instruction pages is available at http://complimet.ca/batl. Users with result tables generated through other acquisition packages can simply use the program by downloading the sample data and formatting their training and test datasets accordingly.

## 3. Results and discussion

BATL was trained on the sphingolipid and glycerophosphocholine training sets with unassigned identity weights *w*_*iu*_ learned by 10-fold cross validation and a precursor/product ion tolerance of 0.5 m/z units. Models were constructed from every subset of features presented in Table 1. These features described peak retention times, intensities, and shapes and can be calculated from the standard outputs of all targeted lipidomic peak-picking software programs (e.g.., MultiQuant, version 3.02, SCIEX). To train BATL, labelled validated datasets were used as training sets by adding an additional column “Lipid_identifier.” This identifier can be any standardized character string used by a laboratory to annotate lipid identity. To calculate BATL-specific peak features (Relative RT, Subtracted RT, Relative Area, Relative Height), an internal standard must be specified by the user and can be identified in the GUI. This internal standard must be present in all samples and all datasets (training and test). For each model, lipid identities were assigned to peaks in the cross validation or holdout sets using either the MAP, constrained MAP, or MWBM decision rules. Two peak identification algorithms, retention time mean and retention time window, were also devised as benchmarks recapitulating manual curation performed on-the-fly by users using MultiQuant to target desired peaks. The retention time mean approach assigned peaks to the single lipid identity in the training set with the closest mean retention time. The retention time window approach computed a retention time range for each lipid identity based on their minimum and maximum observed retention times in the training set. Lipid identities were only assigned to peaks whose retention times unambiguously fell within the window of a single lipid species.

**Table 1.**
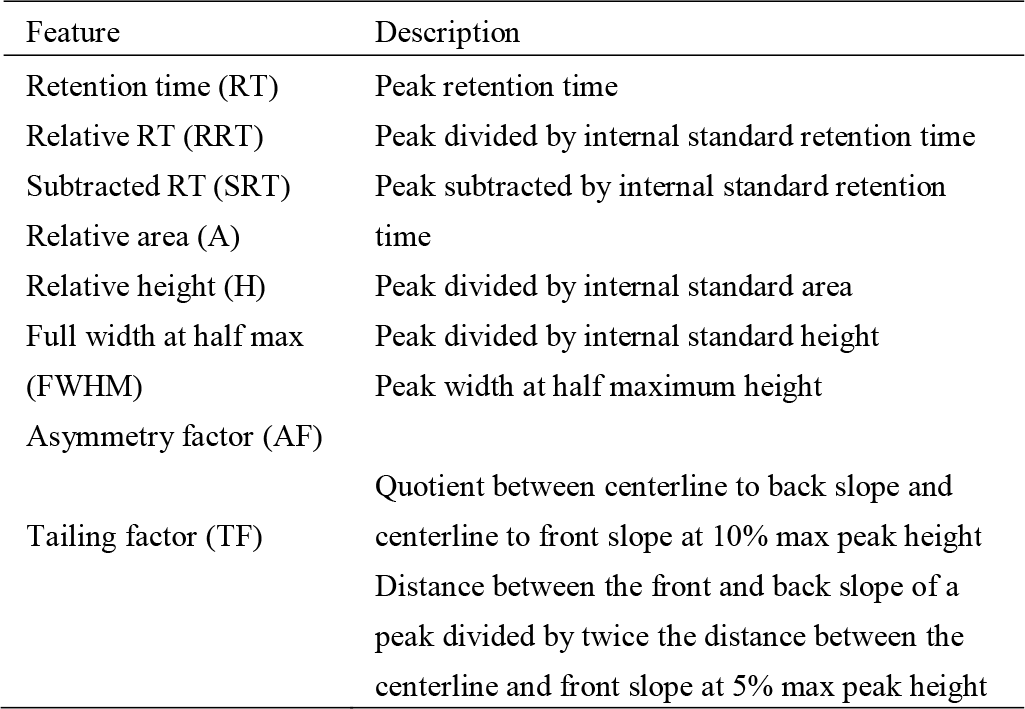
Specified SRM peak features for naïve Bayes model.

To identify the best decision rule, cross validation accuracies were compared between BATL models trained using retention time only but differing in decision rule. For comparison, the accuracies of the two retention time window/mean matching algorithms were included to benchmark the BATL models where Figure 3a) shows over 95% accuracies on the sphingolipid dataset using any method except the retention time window approach. As peaks were only assigned if they fell within the retention time window of a single lipid identity, this method incurred a 10% unassignment rate on the sphingolipid dataset which was two orders of magnitude greater than any of the BATL models (see Supplementary Figure S1a-c).

**Fig. 3.**
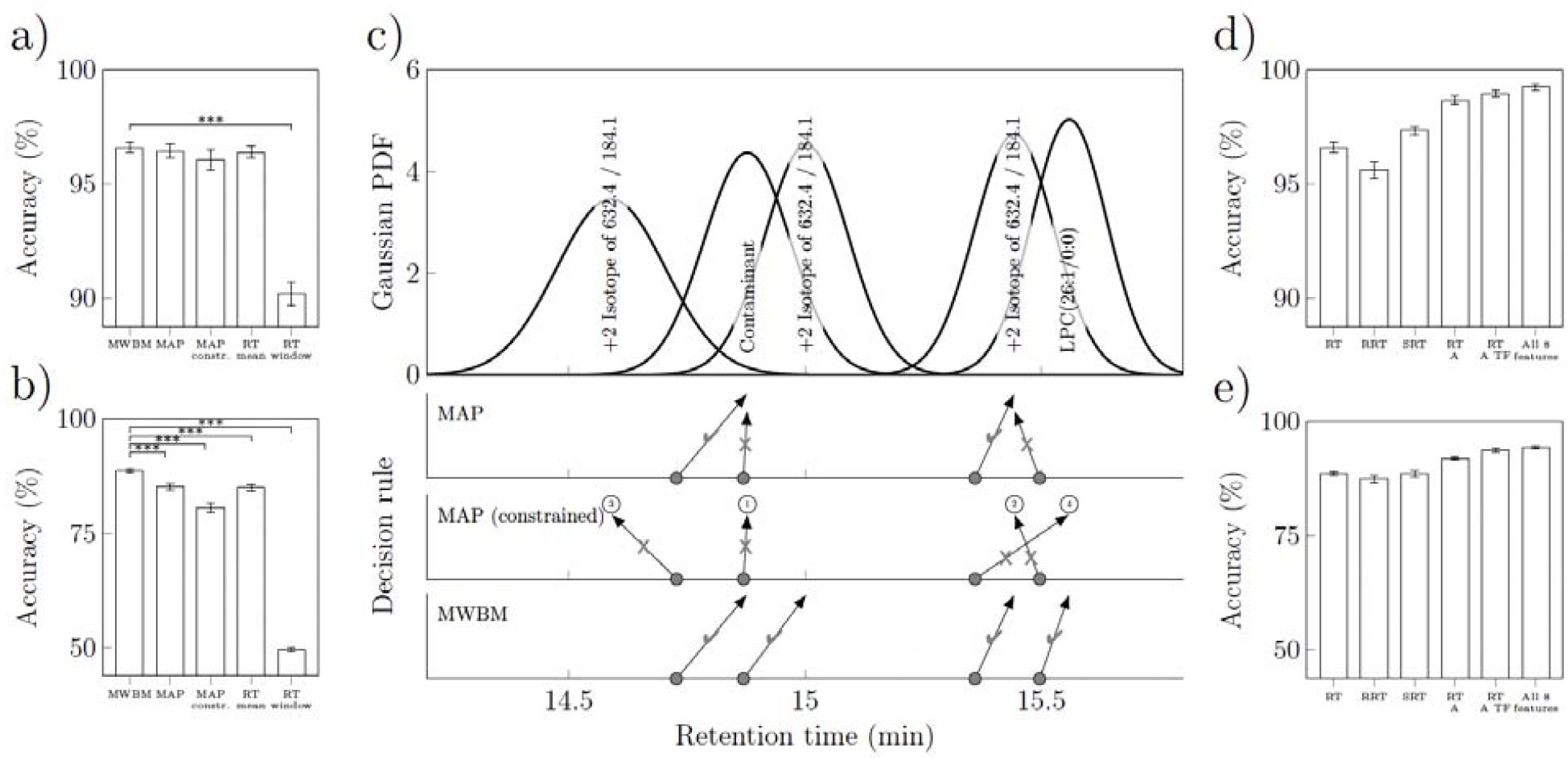
Classifier performance on 10-fold cross validation sphingolipid and glycerophosphocholine datasets. 95% confidence intervals shown in panels a-b, d-e). a-b) Data represent mean accuracies of BATL models trained on retention time with each decision rule and retention time mean/window matching algorithms for a) sphingolipids or b) glycerophosphocholines b) *(***Q* < 0.001, *t*-test adjusted with the Benjamini- Hochberg method of all models against the MWBM decision rule). c) Lipid assignment differences between MAP, constrained MAP, and MWBM decision rules during cross validation and trained using retention time. In the top panel, data represent the Gaussian likelihoods of five glycerophosphocholine isomers based on the retention time feature. The rows of grey dots indicate the retention times of four peaks from the same sample in the validation set. Each row indicates the outcome of the three decision rules. Arrows indicate the lipid assignments; checkmarks indicate correct assignments; Xs indicate incorrect assignments. The numbers for constrained MAP indicate the order of peak assignments. d-e) Data represent mean accuracies of the BATL models using MWBM decision rule trained on several features and feature combinations for d) sphingolipids or e) glycerophosphocholines. The feature name codes are described in Table 1.

Similar accuracies were observed across the naïve mean approach and three BATL models, given the relatively low isobaric complexity of the sphingolipid dataset. Only 56.3% of the peaks in the validation sets matched between two to four lipid isobars at the same transition in the training set (Supplementary Table S1-2). Thus, a large proportion of peaks were guaranteed to match to their corresponding lipid identity. When single lipid targets were detected at a transition, the MWBM decision rule assigned the same peak identities as the MAP or constrained MAP decision rule.

The strengths of different BATL models emerged when classifying the more complex glycerophosphocholine dataset in Figure 3b), where 95.3% of all peaks in the validation datasets were present in transitions that contained at least two and up to eight unique lipid isomers (Supplementary Table S3-4). The BATL model, using the MWBM decision rule, achieved an 88.7% accuracy and significantly outperformed every other method (Supplementary Figure S1d-f). Performances were recapitulated when analyzing the holdout sets (Supplementary Information S1), and similar increases in accuracy were also observed when comparing decision rules of models trained using other feature subsets (See Supplementary Figures S2 and S3).

To understand why the MWBM decision rule outperformed the other methods, retention time likelihoods were assessed for the glycerophosphocholine cross validation analyses. Figure 3c) shows the Gaussian likelihoods of five glycerophosphocholine isomers based on the retention time feature. When peak retention times were close together, both the naïve mean approach and MAP decision rule assigned multiple peaks to the same lipid identities. While the constrained MAP decision rule conceptually improved on the MAP decision rule, accuracies were significantly worse on the glycerophosphocholine dataset. Constrained MAP assigned lipid identities by ranked order of posterior probability. These rankings are denoted in Figure 3c) by the ordinal numbers above the assignment arrows. However, interestingly, the most correct peak assignment was not necessarily the one with the greatest posterior probability. As discussed in Figure 1, retention time shifts can cause peak retention times in one sample to misalign to different peaks present in another sample. A similar problem arises when computing the likelihoods of peaks in samples experiencing retention time shifts. Variations in retention time altered the posterior rankings, increasing the likelihood of an incorrect-versus-correct peak-lipid assignment. Thus, once one lipid identity was incorrectly assigned to a given peak, subsequent peaks with similar retention times were misclassified (or not assigned an identity). In contrast, the best performing MWBM decision rule resolved these twotypes of misidentifications.

While retention time is the most common feature for peak identification, it is not the only lipid-specific peak feature or necessarily the most discriminative one. Figure 3d-e) show cross validation accuracies of selected BATL models using the MWBM decision rule trained using different retention time, intensity, and shape features. Across both the less complex sphingolipid and more complex glycerophosphocholine datasets, additional features describing peak intensity and shape increased classification accuracies and identification rates, while decreasing unassignment rates (Supplementary Figure S4). When comparing models trained on the best subset of N features, the use of all eight features consistently resulted in the best identification and unassignment rates on the holdout sets (Supplementary Figure S5). When adding statistically dependent features to the model, diminishing performance returns were observed, although identification and unassignment rates remained equal to or greater than less complex models on the holdout sets. Of the three retention time features explored, models trained using subtracted retention time performed equal to or significantly greater than those trained using regular retention time. Notably, this method of accounting for variations in LC retention time has not been reported in literature. Software programs such as MultiQuant can report both regular and relative retention times if an internal standard is specified. Although relative retention time is designed to correct against retention time shifts, this method of normalization was sometimes found to induce retention time shifts when no systematic retention time differences were observed across samples (i.e., only transient component level variation was detected). A comparison of models trained on each feature using the MWBM decision rule revealed equal to or significantly worse identification rates between relative-versus-regular retention time (Supplementary Figure S6). Overall, models trained using retention time features significantly outperformed peak intensity and shape features which were the least discriminative, while combinations of multiple features outperformed models focusing on single feature characteristics.

To ensure BATL can be used across platforms, researchers are required to develop their own curated training sets specific to their LC methodologies. Limitations of BATL are that the annotations returned by BATL depend on the accuracy of the identifications assigned in the training set and on the size of the training dataset. Supplementary Figure S7 shows the performance of BATL on the holdout sets when trained on 10% increments of the sphingolipid or glycerophosphocholine training sets. Models were trained using the best single feature or all eight features and every 10% increment corresponded to 22 sphingolipid or 24 glycerophosphocholine samples. Whether trained on the less complex sphingolipid dataset or the more complex glycerophosphocholine dataset, identification rates decreased by less than 1% and unassignment rates remained under 5% when training on 10% of samples. These data demonstrate that only a small rigorously validated training set (i.e., 22-24 samples) is required to train the naïve Bayes model for accurate peak identification.

Benchmarking BATL against other state-of-the-art methods for peak classification is challenging because BATL assigns lipid identities to a list of curated SRM peaks provided by the user, as is the nature of a targeted lipidomic approach, while vendor-specific (e.g., LipidSearch, MultiQuant) and free programs (e.g. METLIN-MRM) pick peaks automatedly and output the pre-assigned targeted identities assuming peak-picking accuracy. As a result, for all programs except MRMPROBS, it is not possible to separate peak detection accuracy from peak identification accuracy. Indeed, this is one of the problems BATL seeks to address. BATL was thus benchmarked against MRMPROBS (Tsugawa, et al., 2013). A notable shortcoming of MRMPROBS, however, is that the number of lipids in the training set cannot exceed the number of lipid targets in the SRM method, meaning that MRMPROBS can only compare identical acquisition methods and cannot annotate a peak as “unassigned” or indicate a new isobar has been selected not already present in the training set. It was thus impossible to apply MRMPROBS to the sphingolipid or glycerophosphocholine holdout sets as they contained different numbers of isobars at a given transition in the training set. This problem was overcome by applying MRMPROBS to multiple training sets containing all combinations of lipid isobars, not exceeding the number of sample peaks. In practice, however, MRMPROBS cannot be used to compare matrices wherein different numbers of isobars are present and a user seeks to annotate which lipids are corresponding between two tissues. To compare, BATL and MRMPROBS, we used a technical replicate dataset which applied the exact same SRM method to monitor sphingolipid species present in 33 replicate runs of two human plasma samples. Thus, both training and testing sets contained the same number of lipids *de facto*. For this analysis, 75% of the samples in the dataset were used to train MRMPROBS and BATL. To construct the MRMPROBS training set, the mean retention times of each lipid were computed from the training set, the logistic regression probability threshold was set to 70%, and the retention time deviation parameter was empirically computed following the MRMPROBS guidelines (Tsugawa, et al., 2013). On the remaining 25% of the technical replicate holdout set, 8.9% of all peaks were not detected by MRMPROBS using a fifteen second retention time window to account for peak detection differences between MRMPROBS and MultiQuant which was used to pick peaks in the longitudinal dataset. Excluding the 8.9% undetected peaks, MRMPROBS achieved a 94.5% identification rate and 4.7% unassignment rate, while BATL, trained using all eight features, achieved 100% identification rate and 0.02% unassignment rate.

## 4. Conclusion

We present here a targeted lipidomics classifier BATL which uses a naïve Bayes model and MWBM decision rule to simultaneously assign lipid annotations to all SRM or MRM peaks in a sample. Using sphingolipid and glycerophosphocholine SRM datasets, BATL was validated on holdout sets with accuracies of 95% or greater when trained using all eight features. As a simple probabilistic classifier, identification and assignment rates remained stable when BATL was trained on as few as 22-24 samples. Lastly, BATL was benchmarked against a retention time window and mean matching approach, comparable to many peak identification programs as well as to the MRMPROBS software program. BATL correctly identified more peaks than either approach with lower unassignment rates and no limitations regarding the number of lipids labelled in the training set nor number of transitions present in the test sets.

In summary, we emphasize that BATL is trainable on any continuous feature and applicable to targeted lipidomics data from any vendor or LC-ESI-MS/MS platform with proper data input formatting. Learning the posterior probability cutoffs is simple to compute based on the naïve Bayes assumption, taking less than ten minutes to train each model on the sphingolipid and glycerophosphocholine training sets analyzed in this study using an Intel i5-8350U mobile processor. To facilitate user experience, we provide BATL at http://complimet.ca/batl wherein instructions and user prompts ensure a fluid training and test pipeline. The sample datasets take less than 10 min to process online with respect to building the BATL model. Larger training datasets of > 100 samples can take up to 20 min to generate the BATL statistical model. Further reductions in classifier training times on larger training sets are now possible with support for parallel processing and are under development. Multiple training datasets can be compressed as .zip files and uploaded as single .zip file. Users are invited to use the sample biological data wherein the minimum number of training datasets (n=24) are provided as two files that can be .zip for upload and used to annotate an n=12 test dataset. While BATL was validated using SRM data, the program is flexible to operate on other targeted lipidomics data acquisition modes that output lists of peaks detected at precursor and product ion pairs including MRM, neutral loss, precursor ion scan, and product ion scan acquisition modes with caveat that file format must be identical to the sample datasets provided. As BATL does not operate on raw MS data, users can continue using their preferred software program to select their lipid targets and conveniently output peak text files into BATL for identification.

## 5. Funding

This work was supported in part by operating grants RGPIN-2019-06796 to SALB and RGPIN-2019-06604 to TJP from the Natural Sciences and Engineering Research Council of Canada (NSERC), AI-4D-102-3 to SALB and MCC from the National Research Council AI for Design Challenge Program, as well as an NSERC CREATE Matrix Metabolomics Training grant to SALB and TJP. TTN and JGC received an NSERC CREATE Matrix Metabolomics Scholarship. JGC was further supported by an NSERC Alexander Graham Bell Canada Graduate Scholarship and an Ontario Graduate Scholarship. IA received a Parkinson Research Consortium Crabtree Family Fellowship. MGS was supported by the Department of Medicine and the Sam and Uttra Bhargava Family.

### Conflict of Interest

*none declared*.

## Notes

https://complimet.ca/batl/

### Competing Interest Statement

The authors have declared no competing interest.

### Summary of Updates

BATL software is now freely accessible online at https://complimet.ca/batl/

