## Supplementary material for "BATL: Bayesian annotations for targeted lipidomics": Chitpin et al Supplementary Data (from another version of this preprint: version 2, 2021-09-24)

### BATL: Bayesian annotations for targeted lipidomics Supplementary information

#### 1. Experimental protocols

Human plasma, collected and prepared in $K_2EDTA$ Lavender BD Hemogard tubes (#367863), was extracted from cognitively normal persons, and patients suffering from Alzheimer's Disease, Mild Cognitive Impairment, Dementia with Lewy Bodies, or Parkinson's Disease (n=321 participants) or persons positive or negative for SARS-CoV-2 (n=36 participants). Consent was obtained in accordance with the Ottawa Hospital Research Institute Research Ethics Committee and in agreement with the National Institutes of Health and NINDS and the Icahn School of Medicine at Mount Sinai Research Ethics Committee. Use of human samples for this study was approved ty the University of Ottawa Ethics Review Board, certificate H-06-20-5864 and H-02-19-3400. Murine plasma and brain, specifically hippocampus and temporal cortex, extracts were generated as part of an ongoing genotype and intervention study of lipid metabolism in n=180 wildtype and N5 TgCRND8 mice, a sexually dimorphic mouse model of Alzheimer's Disease (Granger *et al.,* 2016). All procedures were approved by the Animal Care Committee of the University of Ottawa and performed in accordance with the ethical guidelines for experimentation of the Canada Council for Animal Care. Lipid extraction methodology was as described in detail in Xu *et al.* (2013).

Briefly, brain tissue was homogenized in 4 mL acidified methanol (A412P-4; Fisher, Nepean, ON, Canada) containing 2% acetic acid (351271-212; Fisher, Nepean, ON, Canada) using a tissue tearer (985370; BioSpec, Bartlesville, OK, USA). For plasma samples, 100 µl of human plasma or 50 µl of murine plasma were added directly to 4 mL methanol containing 2% acetic acid. Internal standards, 90.7 ng PC(13:0/0:0) [LM1600], 99.54 ng PC(12:0/13:0) [LM1000], 249 ng PE(12:0/13:0) [LM1100], 249 ng PS(12:0/13:0) [LM1300], 133.7 ng Cer(d18:1/16:0-D31) [868516], 133.7 ng GlcCer(d18:1/8:0) [860540], 133.7 ng GalCer(d18:1/8:0) [860538], and 75 ng SM(d18:1/18:1-d9) [860740] were added at time of extraction. All standards were from Avanti Polar Lipids, Alabaster, AL, USA. Sodium acetate (0.1 M; S-2889; Sigma-Aldrich, Oakville, ON, Canada) and chloroform (C298-500; Fisher) were added sequentially to each sample to a final ratio of 2:1.9:1.6 (acidified methanol/chloroform/sodium acetate). Samples were vortexed, incubated on ice for 15 minutes, and centrifuged for 5 minutes at 600 x g at 4°C. The organic phase was collected, and the aqueous phase back-extracted 3 times using chloroform. Each organic phase was combined and dried under a steady stream of nitrogen. Lipids were re-solubilized in 300 µl of anhydrous ethanol (P016EAAN; Commercial Alcohols, Toronto, ON, Canada) and stored under nitrogen gas at -80°C in amber vials (C779100AW; BioLynx, Brockville, ON, Canada).

HPLC was performed on an Agilent 1290 Infinity LC system, equipped by a binary pump, with an autosampler maintained at 4°C. Aqueous mobile phase (Solvent A) contained 0.1% formic acid and 10 mM ammonium acetate. Organic mobile phase (Solvent B) contained acetonitrile/isopropanol (5:2 v/v) with 0.1% formic acid and 10 mM ammonium acetate. Reversed-phase liquid chromatography for sphingolipid assessment was performed on a 100 mm x 250 µm (inner diameter) capillary column packed with either ReproSil-Pur 120 C8 (particle size of 3 µm and pore size of 120 Å or ReproSil-Pur 200 C18 (particle size of 3 µm and pore size of 200 Å, Dr. A. Maisch, Ammerbruch, Germany) for glycerophosphocholine assessment. For the GPC analysis, five µl of sample were mixed with 5 µl of an internal standard mixture consisting of PC(O-16:0-d4/0:0) [2.5 ng, 360906; Cayman Ann Arbor, MI, USA], PC(O-18:0-d4/0:0) [2.5 ng, 10010228; Cayman], PC(O-16:0-d4/2:0) [1.25 ng, 360900; Cayman], and PC(O-18:0-d4/2:0) [1.25 ng, 10010229; Cayman] in EtOH, PC(15:0/18:1-d7) [1.25 ng, 791637; Avanti Polar Lipids], PE(15:0/18:1-d7) [1.25 ng, 791638; Avanti Polar Lipids], LPC(18:1-d7/0:0) [1.25 ng,791643; Avanti Polar Lipids], LPE(18:1-d7/0:0) [1.25 ng, 791644; Avanti Polar Lipids], and PS(15:0/18:1-d7) [1.25 ng, 791639; Avanti Polar Lipids] and 13.5 µl of Solvent A. For sphingolipid analysis, five µl of sample were mixed with 2.5 µl of EtOH and 16 µl of Solvent A. The LC method operated at a flow rate of 10 µl/min with 3 µl of sample injection for GPC analysis and 5 µl for sphingolipid analysis. The LC gradient used for GPC analysis started at 30% Solvent B, reached 100% Solvent B at 8 minutes, and remained at 100% B for 45 min. At 45 min, composition was returned to 30% Solvent B and the column was regenerated for 15 min. For sphingolipid analysis, the gradient began at 30% Solvent B, was ramped to 100% Solvent B over 5 min, and was maintained for 30 min. The composition was returned to 30% Solvent B at 30 min and maintained for 20 min to regenerate the column. A blank run, wherein 3 µl or 5 µl of Solvent A was injected, followed each sample run.

ESI-MS/MS acquisition and instrument control were performed using Analyst software (version 1.6.2, SCIEX) on a QTRAP 5500 triple quadrupole mass spectrometer equipped with a Turbo V ion source (SCIEX). The ion source operated at 5500 V and 0°C for GPC analysis, 250°C for sphingolipid analysis. Nebulizer/heated gas (GS1/GS2), curtain gas (CUR), and collision gas (CAD) were set to 20/0 psi, 20 psi, and medium, respectively, for GPC analysis. For sphingolipid analysis, the values for these source parameters are 20/20 psi, 20 psi, and medium, respectively. Nitrogen was used as GS1/GS2, curtain and collision gas. Compound parameters (declustering potential, entrance potential, collision energy, and collision cell exit potential) were individually optimized for each transition. All data acquisitions were performed in the positive ion mode. The GPC lipidome monitored the diagnostic product ion at m/z 184.1, indicative of the glycerophosphocholine headgroup, while the sphingolipidome was profiled using the diagnostic product ion 264.3 indicative of the sphingosine backbone. MultiQuant software (version 3.0.2 AB SCIEX) was used for peak picking and processing quantitative SRM data.

The identification of lipid species was performed by employing Information-Dependent-Acquisition Enhanced-Product-Ion-Scan (IDA-EPI) following the quantitative MRM acquisition which served as a survey scan. The IDA method triggered EPI scans following analysis of MRM signals with dynamic background subtraction from the survey scan. The IDA criteria were set to select one to three most intense peaks, and intensity threshold was set to exceed 1000 cps. The EPI experiment operated in the positive mode, scanning mass range from m/z 200-1000 at a scan rate of 10,000 Da/s with dynamic fill in the trap. Two different collision energies were applied, 35 and 50 eV with collision energy spread (CES) of 15 eV to ensure a broad coverage of fragmentation.

#### 2. Author contributions

JGG developed the BATL method, conducted the analyses, and drafted the manuscript. AS developed the GUI interface and online implementation of BATL at http://complimet.ca/batl/. RO, JJT, AMC, MM, MGS, and RSP provided the human samples. TTN, GPT, HX, and IA generated all animal samples, prepared all human and animal samples and generated all of the mass spectrometry data. TJP and SALB guided the analysis of the core BATL program and revised the manuscript. MCC and SALB guided the analysis of the online implementation and revised the manuscript. All authors read, edited, and approved the manuscript.

#### 2. Dataset complexity

**Table 1: Sphingolipid training set isomeric/isobaric complexity.**

| Number of candidate assignments | Detected peaks | Exact mass assignments[1] |
|---|---|---|
| 1 | 13222 | 13222 |
| 2 | 11403 | 704 |
| 3 | 4590 | 94 |
| 4 | 1023 | 0 |

[1]Peak matches the correct lipid target based on Q1 and Q3 m/z alone, regardless of the feature value(s).

**Table 2: Sphingolipid holdout set isomeric/isobaric complexity.**

| Number of candidate assignments | Detected peaks | Exact mass assignments[1] |
|---|---|---|
| 1 | 13069 | 13069 |
| 2 | 11307 | 800 |
| 3 | 4545 | 159 |
| 4 | 1008 | 0 |

[1]Peak matches the correct lipid target based on Q1 and Q3 m/z alone, regardless of the feature value(s).

**Table 3: Glycerophosphocholine training set isomeric/isobaric complexity.**

| Number of candidate assignments | Detected peaks | Exact mass assignments[1] |
|---|---|---|
| 1 | 9492 | 9492 |
| 2 | 16880 | 1034 |
| 3 | 39002 | 0 |
| 4 | 52940 | 173 |
| 5 | 45202 | 0 |
| 6 | 25033 | 0 |
| 7 | 7490 | 0 |
| 8 | 5831 | 0 |

[1]Peak matches the correct lipid target based on Q1 and Q3 m/z alone, regardless of the feature value(s).

**Table 4: Glycerophosphocholine holdout set isomeric/isobaric complexity.**

| Number of candidate assignments | Detected peaks | Exact mass assignments[1] |
|---|---|---|
| 1 | 9474 | 9474 |
| 2 | 16963 | 1050 |
| 3 | 38838 | 0 |
| 4 | 53361 | 181 |
| 5 | 44937 | 0 |
| 6 | 25760 | 0 |
| 7 | 7483 | 0 |
| 8 | 5810 | 0 |

[1]Peak matches the correct lipid target based on Q1 and Q3 m/z alone, regardless of the feature value(s).

**Table 5: Feature codes unless otherwise stated in the following supplementary figure captions.**

| **Feature** | **Abbreviation** |
|---|---|
| Retention time | RT |
| Relative retention time | RRT |
| Subtracted retention time | SRT |
| Relative area | A |
| Relative height | H |
| Full width at half maximum | FWHM |
| Asymmetry factor | AF |
| Tailing factor | TF |

##### 3. Classifier holdout performance between decision rules

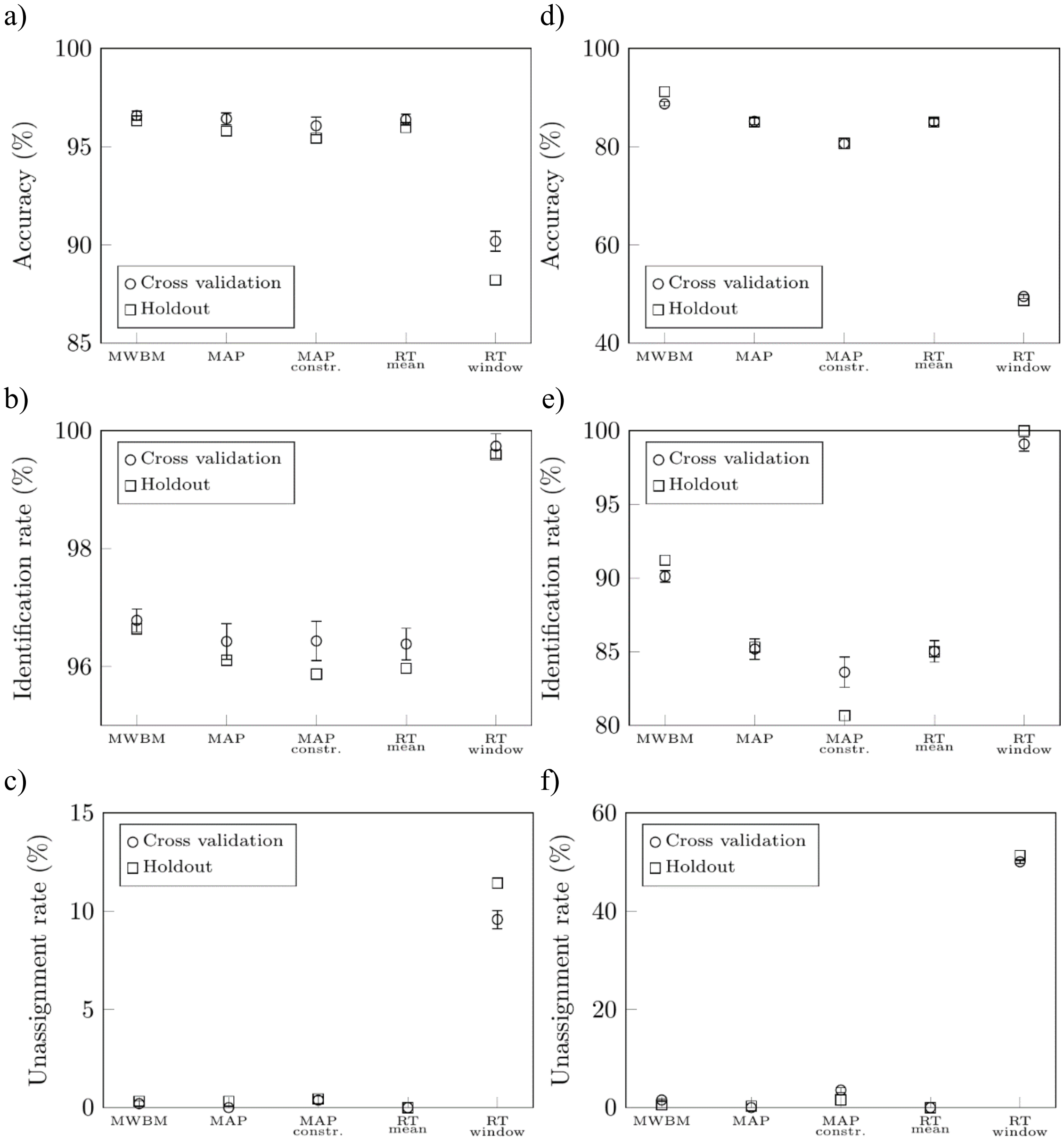

**Figure S1: Detailed classifier performance on holdout and cross validation sets.** For cross-validation results, point markers indicate mean accuracies/identification/unassignment rates and whiskers represent 95% confidence intervals across 10 folds. a-c) Accuracies, identification rates, and unassignment rates for the sphingolipid datasets. d-f) Accuracies, identification rates, and unassignment rates for the glycerophosphocholine datasets.

#### 4. Classifier cross validation performance between decision rules for other feature combinations

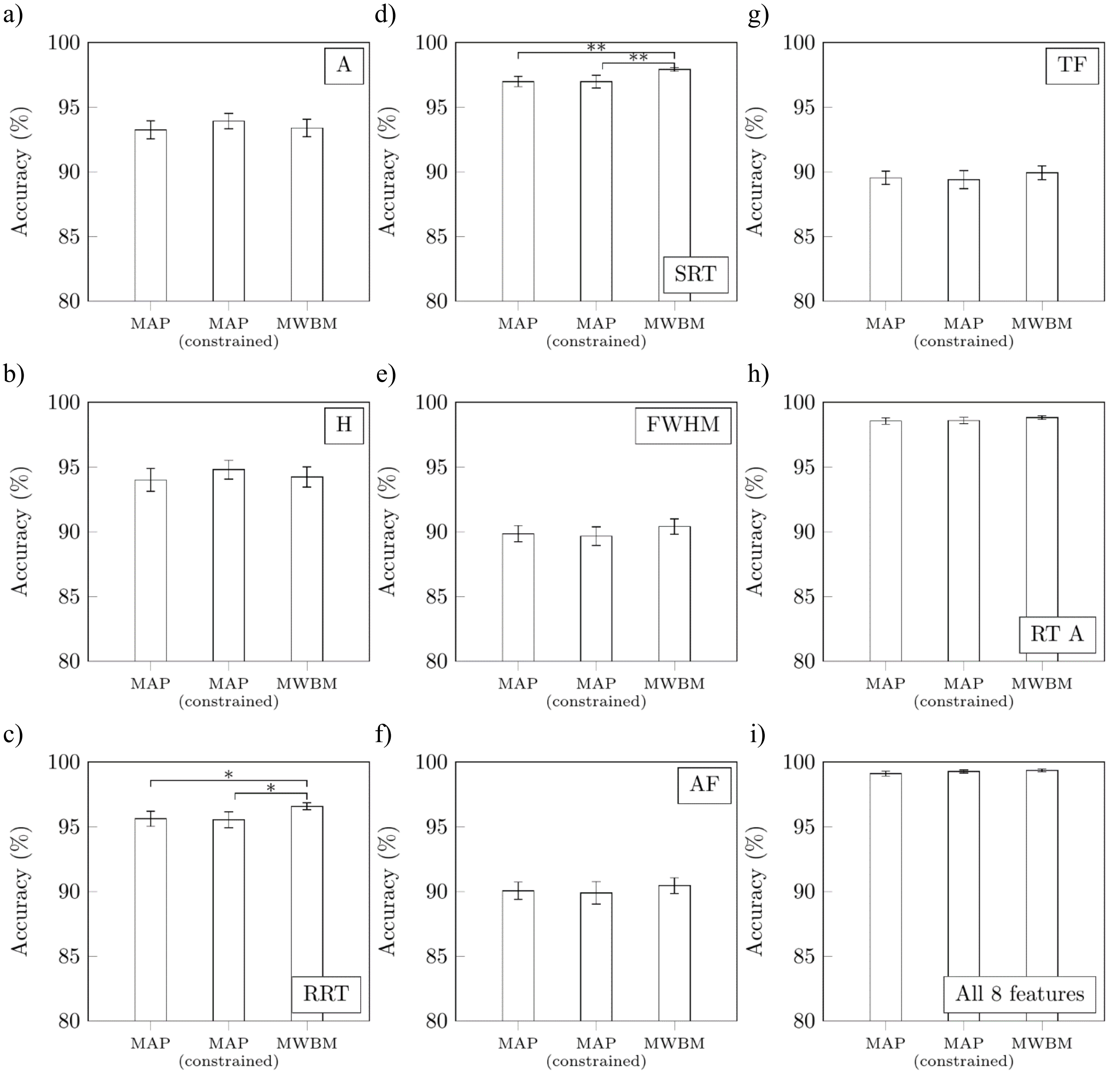

**Figure S2: Classifier performance on 10-fold cross validation sphingolipid datasets.** Data represent mean accuracies of BATL models trained on select feature combinations. 95% confidence intervals shown (**Q* < 0.05, ***Q* < 0.01, ****Q* < 0.001, *t*-test adjusted with the Benjamini-Hochberg method).

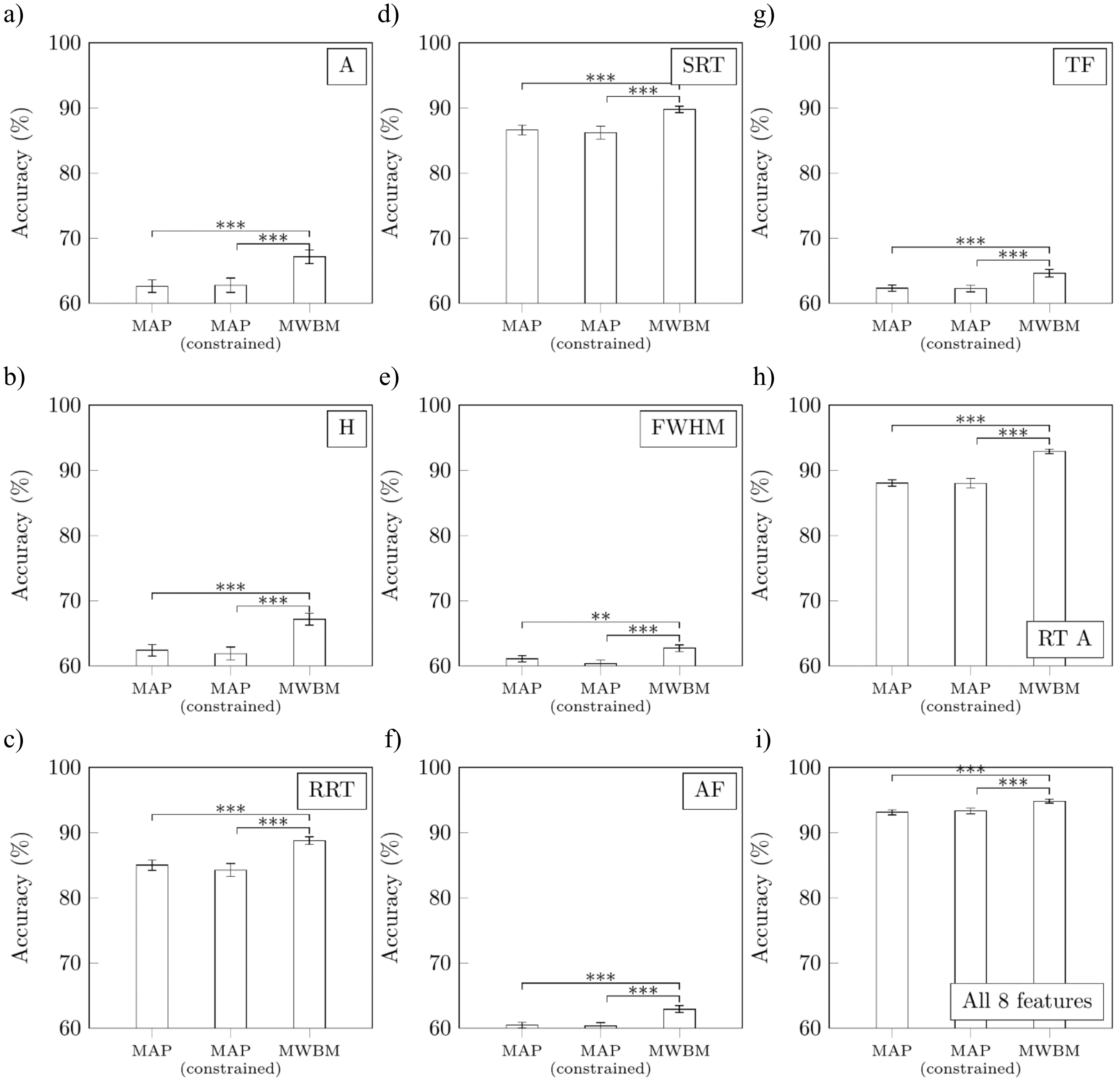

**Figure S3: Classifier performance on 10-fold cross validation glycerophosphocholine datasets trained with select feature combinations.** Data represent mean accuracies of BATL models trained on select feature combinations. 95% confidence intervals shown (*$Q < 0.05$, **$Q < 0.01$, ***$Q < 0.001$, *t*-test adjusted with the Benjamini-Hochberg method).

#### 5. Classifier holdout performance between selected features

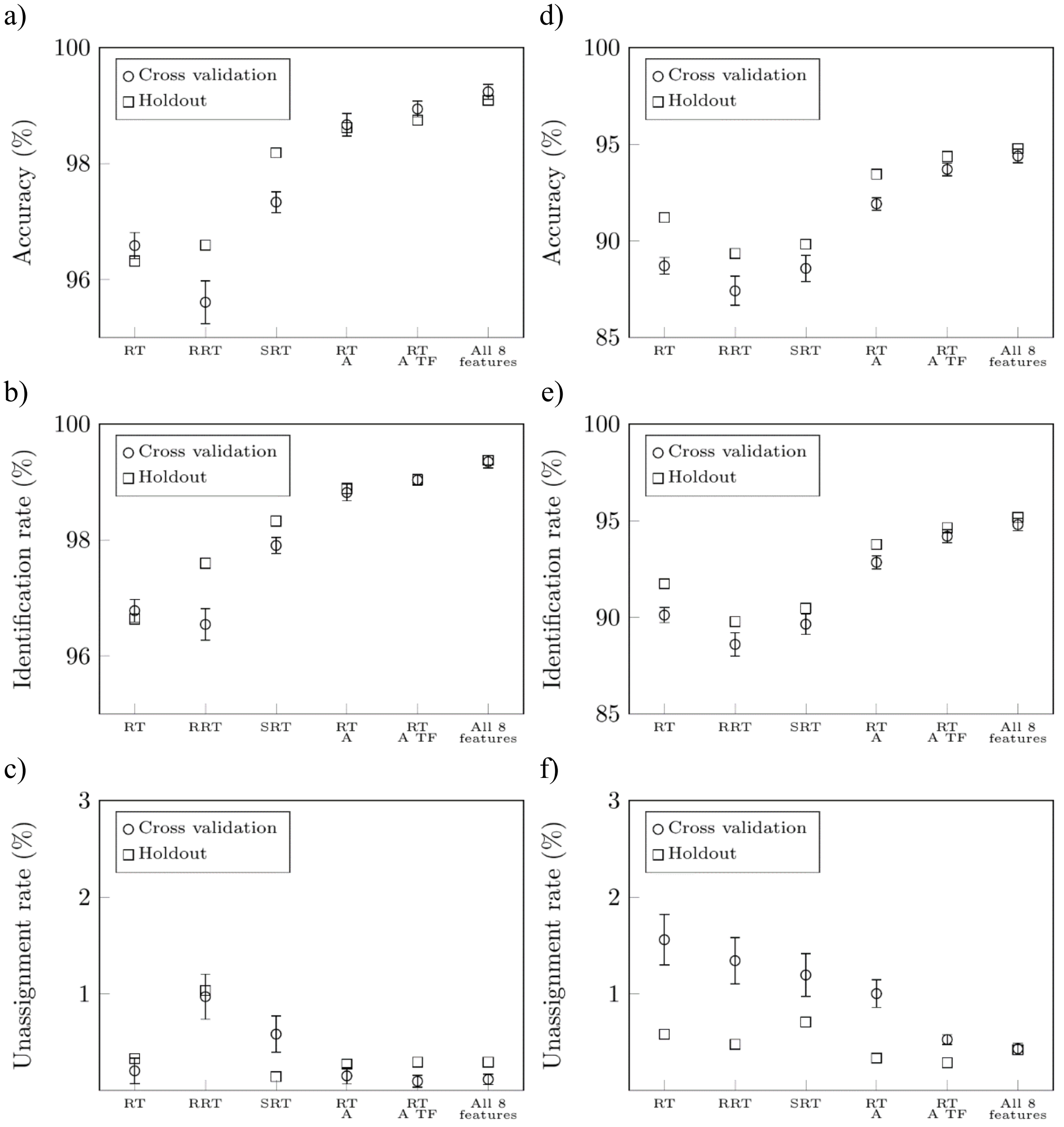

**Figure S4: Classifier performance on holdout and cross validation sets trained with select feature combinations.** For cross-validation results, point markers indicate mean accuracies/identification /unassignment rates and whiskers represent 95% confidence intervals across 10 folds. a-c) Accuracies, identification rates, and unassignment rates for the sphingolipid datasets. d-f) Accuracies, identification rates, and unassignment rates for the glycerophosphocholine datasets.

**6. Classifier performance by best *N* feature combinations**

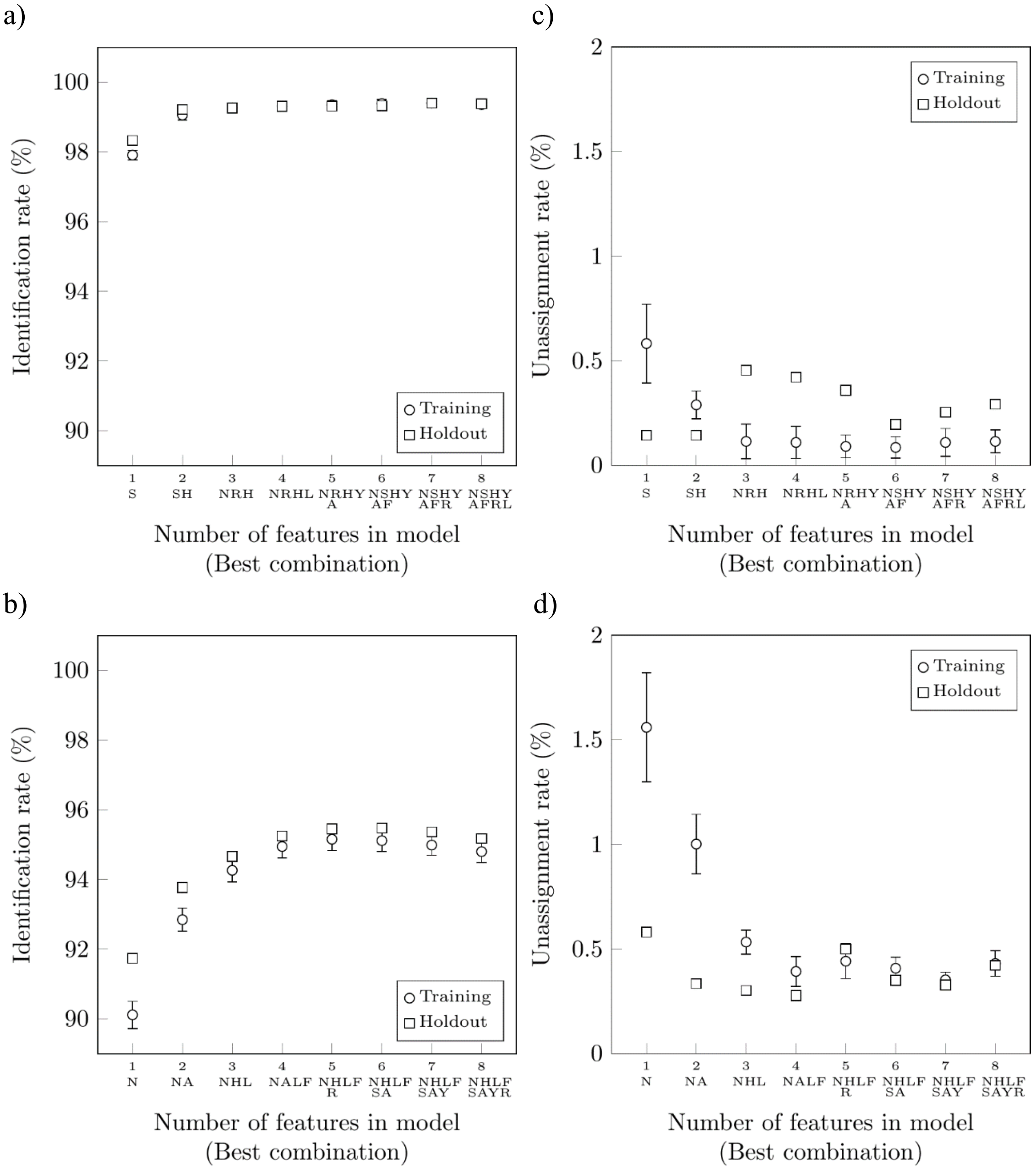

**Figure S5: Models using the MWBM decision rule trained from the best combination of *N* features. X-label feature codes described in Figure S6**. 95% confidence intervals for the 10-fold cross validation datasets shown. a-b) Sphingolipid and glycerophosphocholine dataset identification accuracies. c-d) Sphingolipid and glycerophosphocholine dataset unassignment rates.

##### 7. Classifier performance by single feature

a) b)

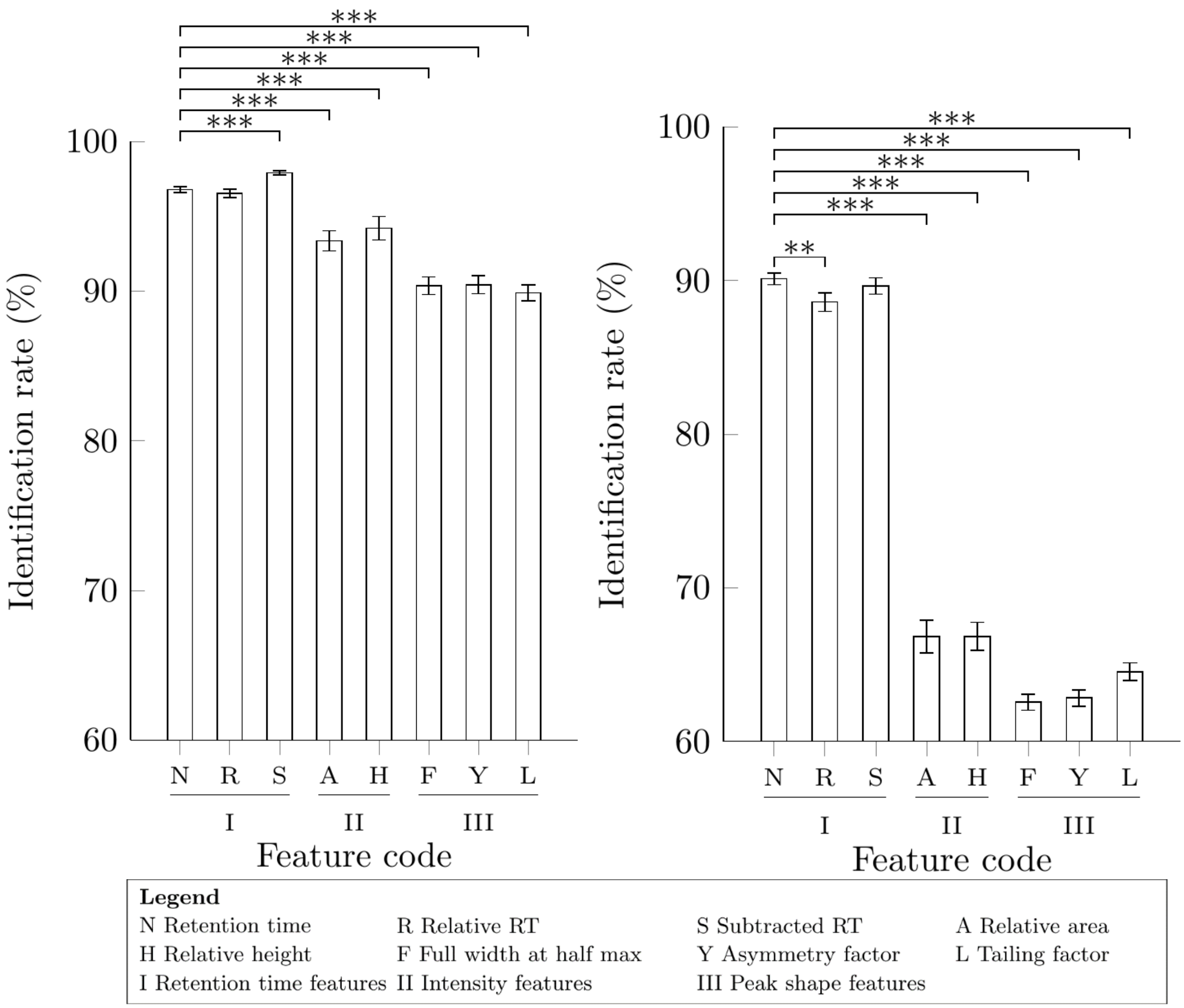

**Figure S6: Comparison of models using the MWBM decision rule trained with retention time (N) versus all other single features.** Ten-fold cross validation identification rates with 95% confidence intervals shown (pairwise *t*-test against the retention time feature adjusted with the Benjamini-Hochberg method $*Q < 0.05$, $**Q < 0.01$, $***Q < 0.001$). a) Sphingolipid cross validation dataset. b) Glycerophosphocholine cross validation dataset.

#### 8. Classifier performance by training set size

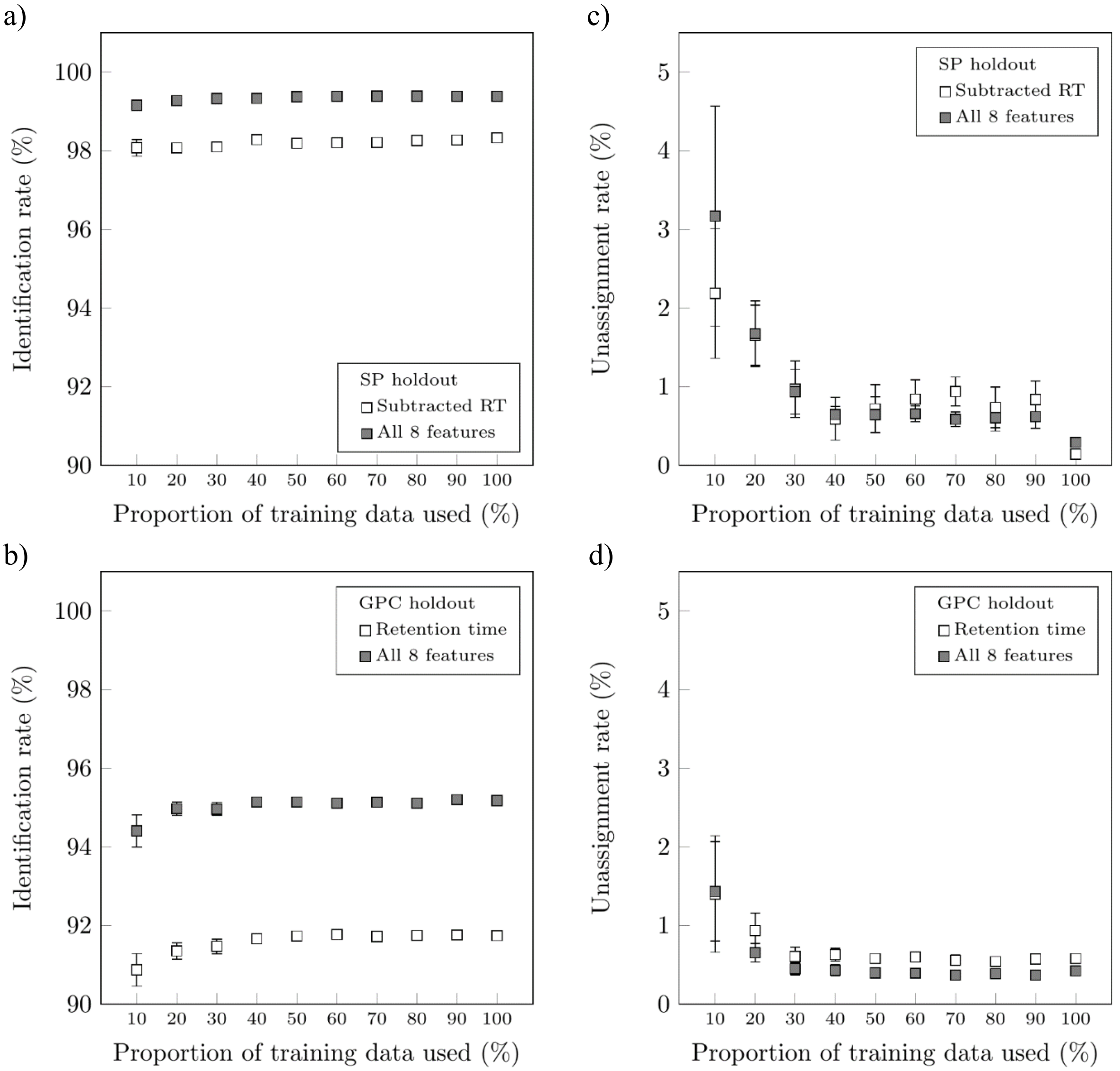

**Figure S7: Classifier performance on holdout datasets with the best single feature or all features resampled ten times for each proportion of training data.** Every 10% increment corresponds to 22 sphingolipid or 25 glycerophosphocholine samples. a-b) Point markers indicate mean identification rates and whiskers represent 95% confidence intervals for the sphingolipid and glycerophoshocholine holdout sets, respectively. c-d) Point markers indicate mean unassignment rates and whiskers represent 95% confidence intervals for the sphingolipid and glycerophosphocholine holdout sets, respectively.
